# Integrated transcriptomics identifies β-cell subpopulations and genetic networks associated with obesity and glycemic control in SM/J mice

**DOI:** 10.1101/2021.07.15.452524

**Authors:** Mario A Miranda, Juan F Macias-Velasco, Heather Schmidt, Heather A Lawson

**Author notes:** Authors contributed equally (Mario A Miranda, Juan F Macias-Velasco). Corresponding author 660 South Euclid Ave, Campus Box 8232, Saint Louis, MO, 63110.

## Abstract

Understanding how heterogeneous β-cell function and stress response impact diabetic etiology is imperative for therapy development. Standard single-cell RNA sequencing analysis illuminates some genetic underpinnings driving heterogeneity, but new strategies are required to capture information lost due to technical limitations. Here, we integrate pancreatic islet single-cell and bulk RNA sequencing data to identify β-cell subpopulations based on gene expression and characterize genetic networks associated with β-cell function in high- and low-fat fed male and female SM/J mice at 20 and 30wks of age. Previous studies have shown that high-fat fed SM/J mice resolve glycemic dysfunction between 20 and 30wks. We identify 4 β-cell subpopulations associated with insulin secretion, hypoxia response, cell polarity, and stress response. Relative proportions of these cells are influenced by age, sex, and diet. Network analysis identifies fatty acid metabolism and β-cell physiology gene expression modules associated with the hyperglycemic-obese state. We identify subtype-specific expression of *Pdyn* and *Fam151a* as candidate regulators of genetic pathways associated with β-cell function in obesity. In sum, this study uses a novel data integration method to explore how β-cells respond to obesity and glycemic stress, helping to define the relationship between β-cell heterogeneity and diabetes, and shedding light on novel genetic pathways with therapeutic potential.

## Introduction

Proper insulin secretion from pancreatic β-cells is required to maintain glycemic control. Obesity initially promotes β-cell expansion, but prolonged glycemic stress and inflammation drive β-cell death and dysfunction, resulting in type 2 diabetes^1–3^. Without sufficient β-cell mass and insulin production, sustained hyperglycemia increases risk for deadly metabolic diseases^4–6^. Currently, transplantation of cadaveric islets is the only method for restoring β-cell mass in diabetes but is severely limited by donor availability and requires lifelong immunosuppressant therapy^7,8^. Differentiating induced pluripotent stem cells into insulin secreting cells may alleviate this bottleneck, but current methods fail to recapitulate fine-tuned glucose sensing and insulin secretion *in vivo*^9,10^. There is urgent need for therapies that improve endogenous β-cell function. This requires understanding how and why β-cells become dysfunctional in obesity.

Improving endogenous β-cell function in diabetes is complicated by cellular heterogeneity, because individual β-cells vary significantly in function, gene expression, protein level, and stress response^11^. Single cell technologies permit interrogation of the molecular underpinnings of heterogeneity, addressing two fundamental questions: Do functionally distinct subpopulations of β-cells exist? Can accounting for heterogeneity improve diabetic therapy? Several research groups have proposed subpopulations based on clustering analysis using single cell RNA sequencing (scRNA-seq), however, there is little agreement among studies^12–15^. High rates of gene dropout and low read depth contribute to these problems, necessitating approaches that improve information capture without losing cell type-specific information^16,17^.

Integrating sc-RNAseq with bulk RNAseq data leverages bulk sequencing’s high read depth, allowing for capture of lowly expressed genes and robust expression analysis. Several tools can integrate sc- and bulk RNA-seq data, focusing on deconvoluting bulk RNAseq data from heterogeneous tissue to account for differences in tissue composition^12,18,19^. These methods estimate and control for cell type abundance but do not identify cell type-specific expression signatures in bulk datasets. Analytical strategies that identify cellular gene expression signatures in bulk RNAseq data allow for robust cell type-specific differential expression analysis and complex network analysis that is currently not feasible in scRNA-seq data.

Here, we characterize gene expression in β-cells from obese SM/J mice, who spontaneously transition from hyperglycemic to normoglycemic with improved β-cell function between 20 and 30 weeks of age^20,21^. We assign functional identities to 4 subpopulations of β-cells using scRNA-seq. These subpopulations vary in proportion among hyperglycemic-obese, normoglycemic-obese, and normoglycemic-lean mice. We identify 316 genes specifically expressed by β-cells and establish a β-cell gene expression profile for each mouse. We leverage this information to focus our analyses of bulk-islet RNAseq and identify β-cell-specific differential expression and gene networks associated with the hyperglycemic-obese, normoglycemic-obese, and normoglycemic-lean states. Two novel potential regulators of β-cell function, *Pdyn* (Prodynorphin) and *Fam151a* (Family with sequence similarity 151 member A), are differentially expressed, highly connected within genetic networks, and primarily expressed by β-cell subpopulations associated with normoglycemia. This analysis demonstrates that integrating scRNAseq with bulk RNAseq is a powerful approach for exploring β-cell heterogeneity and identifying key genes and subpopulations that strongly associate with glycemic state. The genetic networks and β-cell subpopulation signatures we identify have high potential to lead to further research aimed at improving β-cell function in obesity.

## Methods

### Metabolic phenotyping

SM/J mice were obtained from The Jackson Laboratory (Bar Harbor, ME). Mouse colony was maintained at the Washington University School of Medicine and all experiments were approved by the Institutional Animal Care and Use Committee in accordance with the National Institutes of Health guidelines for the care and use of laboratory animals. Mice were weaned onto a high-fat diet (42% kcal from fat; Envigo Teklad TD88137) or isocaloric low-fat diet (15% kcal from fat; Research Diets D12284), as previously described^21^. At 20 or 30 weeks of age, mice were fasted for 4 hours, body weight measured, and blood glucose was measured via glucometer (GLUCOCARD). Mice were injected with sodium pentobarbital, followed by a firm toe pinch to ensure unconsciousness. Blood was collected via cardiac puncture and pancreas was collected. Blood was spun at 6000 rpm at 4°C for 20 minutes to collect plasma. Insulin ELISA (ALPCO 80-INSMR-CH01) was used to measure plasma insulin levels following manufacturer’s instructions.

### Islet isolation and phenotyping

Islets were isolated from pancreas as previously described20 and rested overnight. 5 Islets were equilibrated in KRBH buffer with 2.8 mM glucose for 30 minutes at 37°C, then placed in 150 μl KRBH containing 2.8 mM glucose at 37°C for 45 minutes, then 150 μl KRBH containing 11 mM glucose at 37°C for 45 minutes. Islets were then transferred into 150 μl acid ethanol. Islet content and secretion tubes were stored at −20°C overnight. Experiments were performed in duplicate per individual, and measurements are reported as the average of replicates. Mouse insulin ELISA (ALPCO 80-INSMU-E01) was performed, with the secretion tubes diluted 1:5, and content tubes diluted 1:100. Glucose stimulated insulin secretion was calculated by dividing insulin secretion at 11mM glucose by insulin secretion at 2.8 mM glucose. Basal insulin secretion was calculated by dividing insulin secretion at 2.8 mM glucose by islet insulin content. Total islet protein was measured using Pierce BCA Protein Assay kit (Thermo Scientific) according to manufacturer’s instructions and read at 562 nm on the Synergy H1 Microplate Reader (Biotek). Islet insulin content was calculated by dividing the islet insulin level in the content tubes by total islet protein. All measurements were taken in duplicate, values reported are the average of replicates.

### Single cell RNA sequencing

Single cell RNA sequencing (scRNA-seq) was performed on islets isolated from 15 SM/J mice representing 6 cohorts: 20wk high-fat females (n=3), 20wk high-fat males (n=3), 30wk high-fat females (n=3), 30wk high-fat males (n=2), 20wk low-fat females (n=2), and 20wk low-fat males (n=2). Isolated islets were dissociated into single cell suspensions using Accumax cell/tissue dissociation solution (Innovative Cell Technologies). Libraries were prepped using the Chromium Single Cell 3‘ GEM, Library & Gel Bead Kit v3 (10xGenomics) and sequenced at 2×150 paired end reads using a NovaSeq S4. After sequencing, reads were de-multiplexed and assigned to individual samples. Reads were aligned using 10x Genomics CellRanger (3.1.0) against our custom SM/J reference^22^. Samples that were prepped together were aggregated into batches using CellRanger aggregate. In the R environment (4.0.0), each aggregated batch was run through SoupX (1.5.0)^23^ to estimate and correct for ambient RNA contamination. A contamination fraction of 0.05 was chosen. Removal of *Ins2* ambient RNA shown in **Supplemental Figure 1D–E**. Adjusted counts were imported into Seurat (3.2.2)^24^, where cells were filtered for number of features detected (500-3000), total counts detected (1000-30000), percent mitochondrial genes (0-30), visualized in **Supplemental Figure 1A**. For additional quality control, we excluded cells where nCount was not predictive of nFeature; the predictive error (residual) of a cell had to be within 3 standard deviations of the mean predictive error (~0). Cell counts for samples from one batch shown before and after filtering step shown in **Supplemental Figure 1B–C**. Expression was then normalized in Seurat (normalization.method = LogNormalize), batches were integrated, and clustered using a shared nearest neighbor approach. Using the top 10 principal components of the filtered expression data and a resolution of 0.14, we identified 9 clusters of cells using Clustree (0.4.3)^25^. Cell types were assigned by identifying top over-expressed genes for each cluster relative to all other clusters using a Wilcoxon rank sum test, with an average log-fold-change threshold of >=0.25 and requiring at least 25% of cells express the gene. Identities were assigned by comparing top over-expressed genes for each cluster with known cell-type specific markers for islet cells.

### Identification of β-cell subtypes, differential expression, and composition

All β-cells were subset for further analysis. Using the top 10 principal components of the filtered expression data and a resolution of 0.09 (**Supplemental Figure 1F**), we identified 4 populations of β-cells, labeled Beta 1-4. Top over expressed genes for each population were identified using a Wilcoxon rank sum test, with an average log-fold-change threshold >= 0.1 compared to all other β-cells and an adjusted p-value < 0.01 (Bonferroni), shown in **Table 1**. Subtype-identifier genes were tested for gene set enrichment (see below) to determine presumed function of each population. Using these thresholds (average log-fold-change >= 0.1 and adjusted p-value < 0.01), differential expression was calculated between pairs of β-cell subpopulations (**Table 15-20)**, across all β-cells between 20wk high-fat males and 30wk high-fat males (**Table 21**), across all β-cells between 20wk high-fat males and 20wk low-fat males (**Table 22**), and across Beta 1 cells between 20wk high-fat males and 20wk low-fat males (**Table 23**) using the “MAST” hurdle-model test^26^. Relative proportions of β-cell subpopulations in each cohort was estimated using bootstrapping to calculate 95% confidence intervals by randomly sampling 1,000 cells from each cohort for 100 iterations.

### Identification of β-cell-specific genes

To identify β-cell-specific gene expression signatures in the bulk sequencing data, we assume that for a given gene, the sum of expression in the scRNA-seq data, *Y*_*Total*_, approximates the expression in bulk RNA sequencing data, *Y*_*Bulk*_.

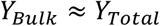

The *Y*_*Total*_ value can be re-written as the sum of expression from all the contributing cell types, where *Y* is the expression from a given cell type, and *N* is the total number of cells of that type.

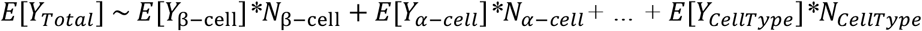

Therefore, the expected relative contribution of β-cells (*Q*_*β-cell*_), to total expression (*Y*_*Total*_), can be written as:

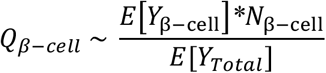

When *Q*_*β-cell*_ is high in all 6 cohorts, we are confident that gene expression in the bulk data is nearly exclusively coming from β-cells. To determine the contribution of β-cells and all other cell types to gene expression, the distribution of normalized expression was assessed using a Cullen and Frey graph using the fitditrplus package (1.1-3)^27^ in R and identified as beta distributed (**Supplemental Figure 2A–B**).

scRNA-seq data is characterized by a buildup of expression values at 0 due to cells that do not express a gene or due to gene-dropout. This distribution required us to employ a beta-hurdle model described by three parameters: Pr(Dropout), α, and β. We treat the probability of attaining an expression value of 0 (Pr(Dropout)) and the distribution of non-zero values separately for a given gene. We fit the expression of each gene in each cell type using a beta-hurdle model optimized by maximum goodness of fit estimation. To fit the first part of the beta-hurdle mode, the Kolmogorov-Smirnov statistic using the ks.test package (4.05)^28^ in R was used to select the optimal α and β shape parameters that best fit the expression distribution of non-zero-expressing cells by minimizing the distance between the cumulative distribution of the real data and theoretical beta model data (Ks) **(Supplemental Figure 2C**). To fit the second part of the beta-hurdle mode, Pr(Dropout) was iterated between 0 and 1, selecting the Pr(Dropout) that minimized Ks (**Supplemental Figure 2D**). In most cases, assuming 100% of cells not expressing the gene was due to gene drop out provided the best fit (**Supplemental Figure 2E**). From the best fit beta-hurdle model, the expected value for each gene within a cell type *E*[*Y*_*CellType*_] can be calculated using the fit α and β shape parameters as:

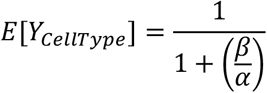

Multiplying the expected value by the total number of cells of that cell type provides the total expression contribution of that cell type. Summing this value across all cell types provides total expression and allows assessment of the contribution of β-cell gene expression to the genes’ total expression. The cell type contribution of all genes in each of the 6 cohorts are shown in **Table 3-8**. We required β-cells to account for >=80% of total gene expression in each of the six cohorts analyzed, resulting in 316 “β-cell-specific genes”, shown in **Table 9**.

### Bulk RNA sequencing

Islets from 32 mice were sequenced: 4 males and 4 females from each diet (high-fat and low-fat) and each age (20wk and 30wk), n=32. Islet RNA was extracted using the RNeasy MinElute Cleanup kit (Qiagen), RNA concentration was measured via Nanodrop and RNA quality/integrity was assessed with a BioAnalyzer (Agilent). Libraries were prepped using the SMARTer cDNA synthesis kit (Takara Bio) and sequenced at 2×150 paired end reads using a NovaSeq S4. After sequencing, reads were de-multiplexed and assigned to individual samples. FASTQ files were trimmed and filtered to remove low quality reads and aligned against a SM/J custom genome using STAR^22,29,30^. Read counts for β-cell-specific genes were normalized via TMM normalization and pairwise differential expression between cohorts was performed using edgeR^31^. Differential expression analysis for all 316 β-cell-specific genes across select cohort comparisons reported in **Table 11**.

### Co-expression network analysis

Weighted Gene Co-Expression Network Analysis (WGCNA) identifies co-expression modules and correlates them with phenotypic traits^32^. Briefly, edgeR-normalized counts for β-cell-specific genes were converted to standard normal, and an adjacency matrix was created from bi-weight mid-correlations calculated between all genes in all individuals and raised to a power β of 8, chosen based on a scale-free topology index above 0.9, to emphasize high correlations^33^. The blockwiseModules function created an unsigned Topological Overlap Measure using the adjacency matrix to identify modules of highly interconnected genes. Eigengenes were calculated as the first principal component for each module, and Pearson’s correlations were calculated between eigengene expression and phenotype to estimate module-trait relationships. Module-trait correlations were considered significant at an FDR-corrected p-value < 0.05. Each gene’s module membership and correlation with metabolic phenotypes are reported in **Table 13**. The adjacency matrix was then used to calculate the connectivity of each gene with other genes within its module (k_within_). Adjacency matrices were subset for each age x diet x sex cohort, used to calculate cohort-specific connectivity for each gene (**Table 14**), then used to calculate differential connectivity between cohorts. We report differential connectivity between 20wk high-fat males and 30wk high-fat males, *HM*, calculated as

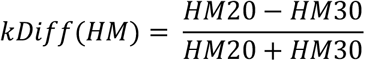

and differential connectivity between 20wk high-fat males and 20wk low-fat males, *M20*, calculated as

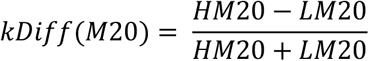

where HM20, HM30, and LM20 are cohort-specific k_within_ values for each gene. This provides each gene in the comparison with a value between −1 and 1, where │kDiff│ > 0.5 is considered differentially connected. Genes with positive differential connectivity are more highly connected in 20wk high-fat males than 30wk high-fat males or 20wk low-fat males, depending on comparison.

### Gene set enrichment analysis

Over-representation analysis (ORA) was performed using the WEB-based Gene Set Analysis Toolkit v2019^34^ on genes overexpressed in individual β-cell subtypes (**Table 2**), β-cell-specific genes (**Table 10**), and genes within blue and brown expression modules (**Table 12**), using all genes expressed in the sc-RNAseq data set as a reference set. Analysis included gene ontologies (biological process, cellular component, molecular function), pathway (KEGG), and phenotype (Mammalian Phenotype Ontology).

## Results

### SM/J islets contain four β-cell subpopulations

We developed an analysis pipeline that integrates sc- and bulk RNA-seq data to characterize the transcriptional changes in β-cells from obese SM/J mice as they improve glycemic control with age (**Figure 1A**). We identified islet cells (α, β, and δ), exocrine cells (acinar and ductal), vascular cells (smooth muscle and endothelial), and immune cells (B cells and macrophages) (**Figure 1B**). Subsequent clustering of β-cells revealed 4 subpopulations, labeled Beta 1-4 (**Figure 1C, Supplemental Figure 1F).** 20wk hyperglycemic high-fat males have a significantly larger Beta 1 population compared to 30wk normoglycemic high-fat males and 20wk low-fat males, at the expense of a diminished Beta 2 population (**Figure 1 D–E**).

**Figure 1.**
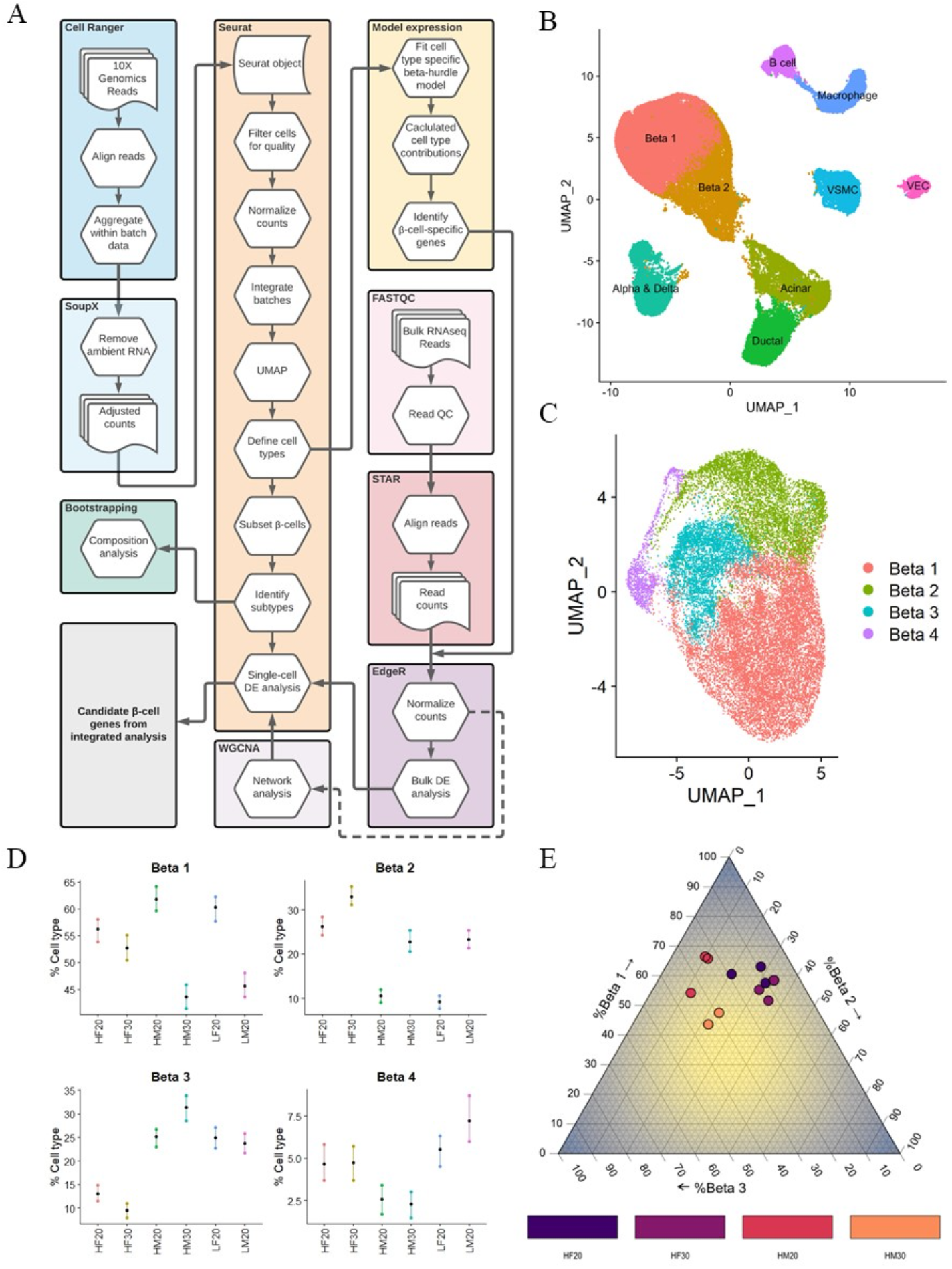
SM/J islets contain four subpopulations of β-cells. **(A)** Data analysis pipeline identifies subpopulations of β-cells and integrates bulk and single cell RNA sequencing data. **(B)** Single cell RNA sequencing UMAP plot of islet tissue. VSMC – vascular smooth muscle cell, VEC – vascular endothelial cell. **(C)** Single cell RNA sequencing UMAP of 4 subpopulations of β-cells. **(D)** Bootstrap analysis quantifies relative proportions of β-cell subpopulations across cohorts. Black dot represents estimated proportion, colored bars illustrate 95% confidence interval. Cell type composition estimates with non-overlapping confidence intervals are significantly different. **(E)** Ternary plot illustrates differences in subtype composition of Beta 1-3 in high-fat SM/J mice. HF20 – 20wk high-fat female, HF30 – 30wk high-fat female, HM20 – 20wk high-fat male, HM30 – 30wk high-fat male, LF20 – 20wk low-fat female, LM20 – 20wk low-fat male.

### β-cell subpopulations have unique expression signatures

To identify genes that are overexpressed in each subpopulation, we performed differential expression analysis between each subpopulation and all other β-cells (**Table 1**). The top ten highest differentially expressed genes within each subtype are visualized in **Figure 2**. Gene enrichment analysis on the overexpressed genes in each subpopulation reveals potential specialization: Beta 1 - insulin secretion, Beta 2 – hypoxia response, Beta 3 – cell polarity, Beta 4 – stress response (**Table 2**).

**Figure 2.**
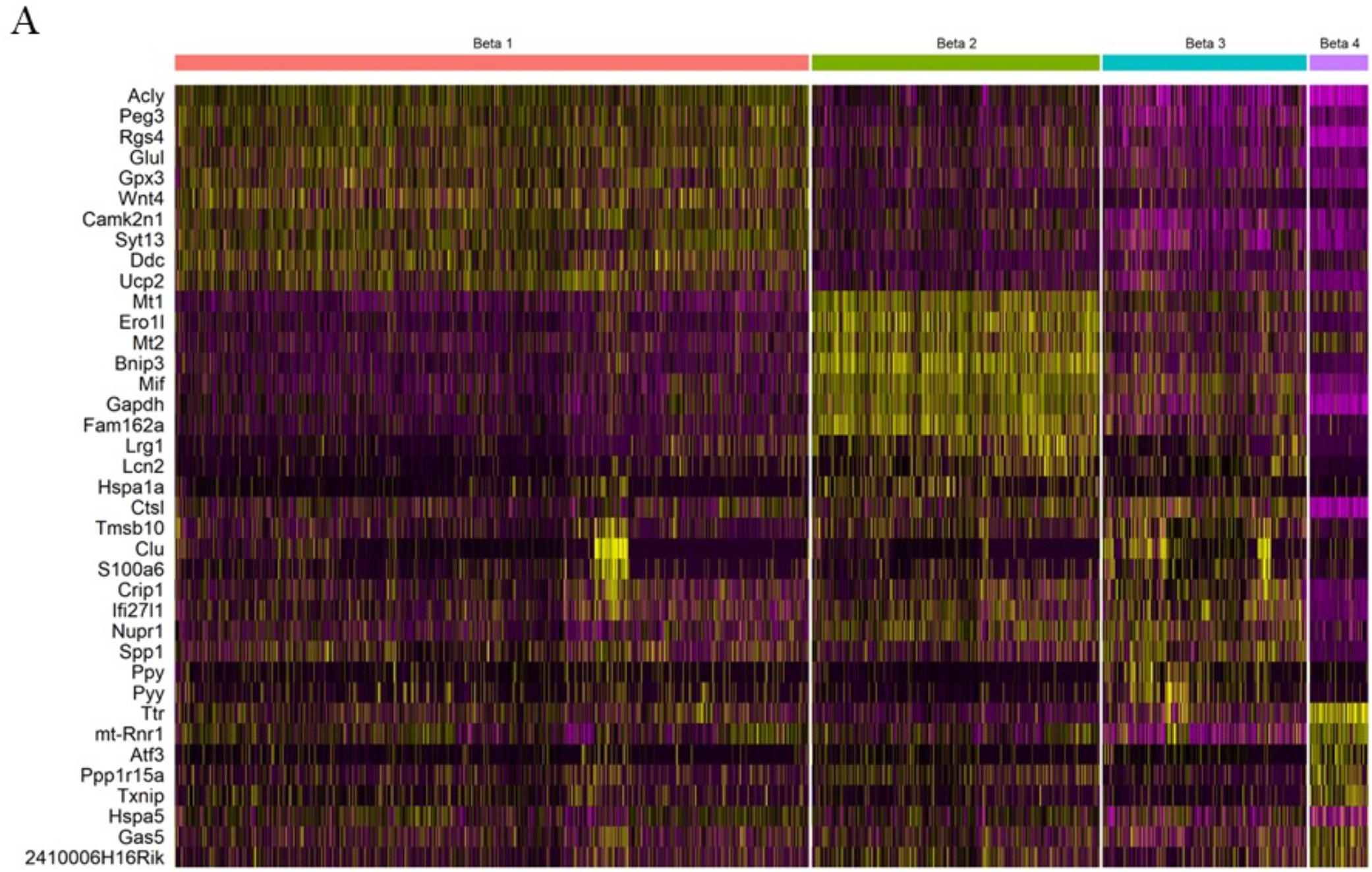
β-cell subpopulations have unique gene expression signatures. **(A)** Heat map for top 10 differentially expressed genes in each β-cell subpopulation. Yellow is highly expressed; purple is lowly expressed.

### SM/J β-cells uniquely express 316 genes

To identify genes primarily expressed by β-cells in scRNA-seq data, we employed a beta hurdle model, which allowed us to estimate the relative contribution of each cell type to total gene expression (**Supplemental Figure 2, Tables 3-8**). To be considered a β-cell-specific gene, we required β-cells to account for at least 80% of total expression within each cohort (**Figure 3A**). We identified 316 β-cell-specific genes (**Table 9**), comprised of genes canonically associated with β-cell identity, including *Ucn3*, *Gcgr*, and *Slc2a2* (**Figure 3B–E**) and genes with unknown function in β-cells (**Figure 3F–G**). Overrepresentation analysis revealed enrichment for terms associated with β-cell function, including mature onset diabetes of the young (MODY) and carbohydrate homeostasis, along with terms related to neuron function including neuroactive ligand-receptor interaction, neuron projection terminus, and axon part (**Table 10**). We then sought to discover if β-cell-specific genes were overexpressed within any of the β-cell subpopulations. We identified 20 β-cell-specific genes overexpressed in Beta 1 cells, far more than would be expected by chance (**Figure 3H**).

**Figure 3.**
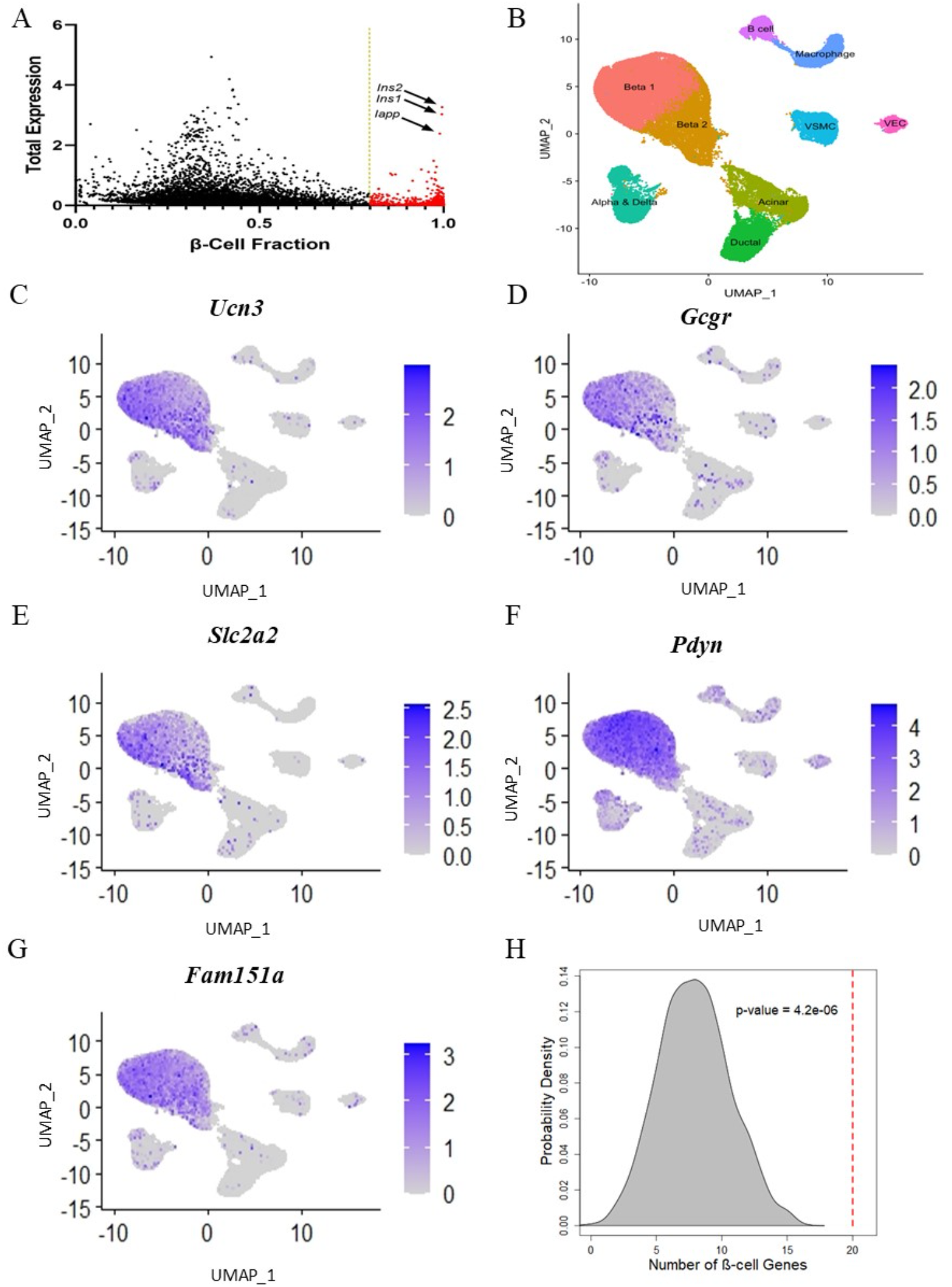
Single cell RNA sequencing identifies β-cell-specific genes. **(A)** Total gene expression and β-cell-specific contribution to gene expression in 20wk low-fat female β-cells. Gold line indicates threshold for β-cell-specific expression. Red dots identify genes that pass the threshold cutoff in this cohort. Arrows identify highly expressed genes associated with β-cell identity. Genes must pass threshold in all cohorts to be considered β-cell-specific in the rest of the analysis. **(B)** UMAP plot of all islet cell types. UMAP plots for β-cell-specific expression of known β-cell markers **(C)** *Ucn3*, **(D)** *Gcgr*, **(E)** *Slc2a2*. UMAP plots for β-cell-specific expression of novel β-cell markers **(F)** *Pdyn*, and **(G)** *Fam151a*. **(H)** Permutation analysis of β-cell-specific genes in Beta 1 overexpressed genes. Distribution shows expected number of β-cell-specific genes in Beta 1 overexpressed gene set based on chance (n= 1000 permutations), red line indicates real number of β-cell-specific genes in Beta 1 overexpressed gene set. P-value indicates probability of number of β-cell-specific genes in Beta 1 overexpressed gene set due to chance.

### β-cell genes are differentially expressed by age, diet, and sex

To robustly characterize β-cell gene expression in SM/J mice, we normalized bulk RNA-seq data from 20- and 30-week high- and low-fat males and females (4 per cohort, n=32) using only the 316 β-cell-specific genes. Principal components analysis on the normalized expression data revealed high-fat males separated from other cohorts (**Figure 4A**). This is consistent with our previous studies showing that high-fat fed SM/J males show a more extreme glycemic response than other cohorts^20,21^. Pairwise comparison revealed 8 differentially expressed genes between 20- and 30wk high-fat males and 2 differentially expressed females between 20- and 30wk high-fat females (**Figure 4B**). 20wk male mice differentially express 104 genes between diets, while 30wk male mice differentially expressed 17. Females differentially expressed a largely consistent set of genes between diets in 20- and 30wk cohorts. Differential expression analysis is reported in **Table 11**. We focused subsequent analyses between 20wk and 30wk high-fat males (**Figure 4C**), and 20wk high and low-fat males (**Figure 4E**). These analyses allow comparison between hyper- and normoglycemic-obese mice, and between hyperglycemic-obese and normoglycemic-lean mice. We highlight two genes with unknown roles in β-cell function, *Pdyn* and *Fam151a*, as differentially expressed between 20- and 30wk high-fat males (**Figure 4D**), and 20wk high and low-fat males (**Figure 4F**), respectively.

**Figure 4.**
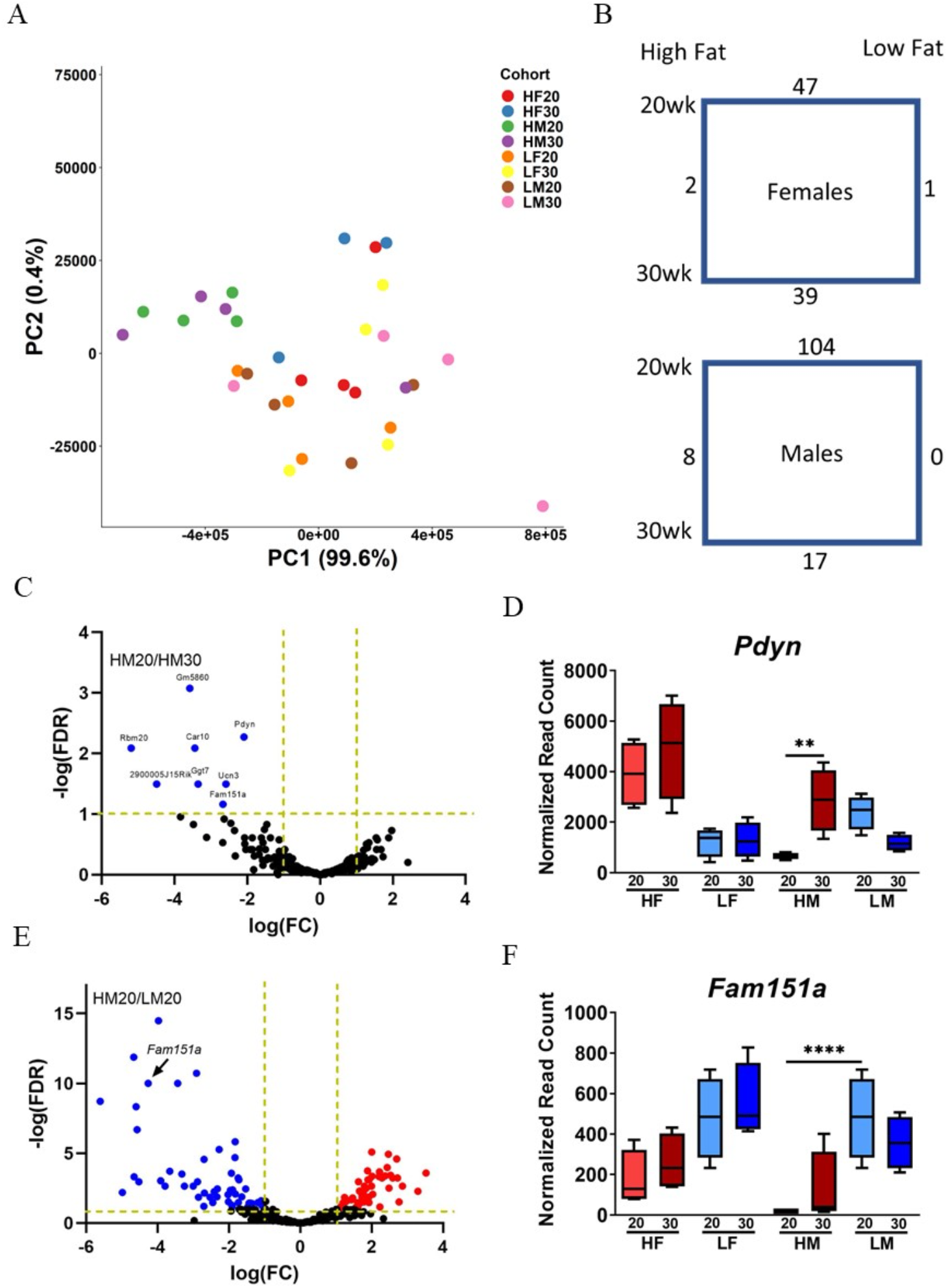
β-cell-specific genes are differentially expressed by age, diet, and sex. **(A)** Principal component analysis of bulk RNA sequencing data normalized with only β-cell-specific genes. **(B)** Number of differentially expressed genes in females and males, across diets and age. **(C)** Differentially expressed genes between 20wk high-fat males and 30wk high-fat males. **(D)** *Pdyn* is differentially expressed in high-fat males. **(E)** Differentially expressed genes between 20wk high-fat males and 20wk low-fat males. **(F)** *Fam151a* is differentially expressed in 20wk males. Blue genes are significantly under-expressed in comparison, red genes are significantly over-expressed. Vertical golden lines indicate threshold for significance based on average log fold change, horizontal line indicates threshold for significance based on FDR corrected p-value. **p-value <0.01, **** p-value < 0.0001 in FDR corrected pairwise comparison. HF20 – 20wk high-fat female, HF30 – 30wk high-fat female, HM20 – 20wk high-fat male, HM30 – 30wk high-fat male, LF20 – 20wk low-fat female, LM20 – 20wk low-fat male.

### β-cell gene expression networks correlate with metabolic traits

We performed network analysis to characterize how groups of genes behaved across age and dietary cohorts. We performed weighted gene co-expression network analysis (WGCNA), which groups similarly co-expressed genes into discreet modules, then correlates these modules with phenotypes^32^. We collected metabolic phenotypes in the bulk RNA-seq mice including body weight, blood glucose level, serum insulin, and islet-specific phenotypes including glucose-stimulated insulin secretion, basal insulin secretion, and islet insulin content (**Supplemental Figure 3**). Performing co-expression analysis identified 9 discreet modules of genes, 6 of which correlated significantly with at least one phenotype (**Table 13, Supplemental Figure 4**). We highlight the blue and brown modules for their correlation with phenotypes relevant to hyperglycemic obese 20wk high-fat mice.

### Blue module associated with fatty acid metabolism correlates with blood glucose level and is altered by age in obese mice

The blue module contains 42 genes, and over-representation analysis reveals enrichment for genes related to fatty acid metabolism (**Table 12**). Blue network eigengene expression correlates with blood glucose level across all mice (**Figure 5A**, **Supplemental Figure 5F**). This correlation is driven by the resolution of hyperglycemia in high-fat mice (**Figure 5A**, **Supplemental Figure 5B**). Strength of individual gene membership within the blue module correlates with strength of correlation to blood glucose levels (**Figure 5B**). The blue network is visualized as the sum of gene-pair correlations (**Supplemental Figure 5D**). This network was then subset into 20wk high-fat males (**Figure 5D**) and 30wk high-fat males (**Figure 5E**). Four genes were identified as differentially connected between networks, although none were individually differentially expressed between 20- and 30wk high-fat males (**Figure 5C, Table 14**).

**Figure 5.**
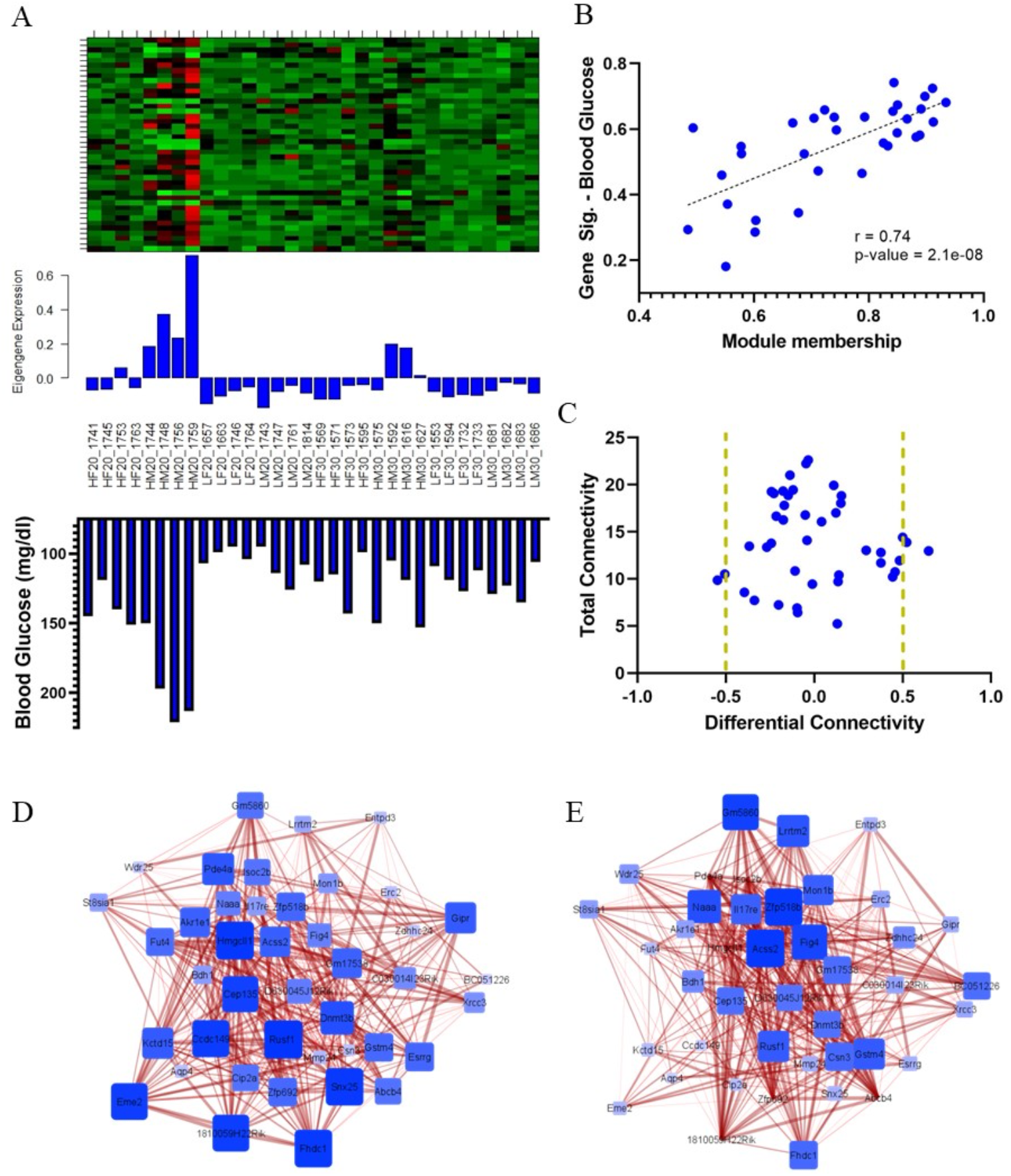
Blue modules network structure is altered by age in high-fat male mice. **(A)** Blue module heatmap, eigengene expression, and blood glucose levels across all individuals. **(B)** Correlation between strength of module membership and blood glucose levels for blue module. **(C)** Total and differential connectivity between blue module genes in 20wk and 30wk high-fat male mice. Vertical golden lines indicate threshold for differential connectivity. **(D)** Blue module network structure in 20wk high-fat males and **(E)** 30wk high-fat males. Size and color of node indicates overall connectivity within the network, thickness of edges indicates strength of correlation between gene pairs.

### Brown module associated with β-cell function correlates with insulin levels and is influenced by diet

The brown module contains 39 genes, and over-representation analysis reveals enrichment for genes related to abnormal β-cell physiology (**Table 12**). Brown network eigengene expression negatively correlates with serum insulin levels across all mice (**Figure 6A, Supplemental Figure 5E**). This correlation is driven in part by diet, where high-fat mice have high levels of circulating insulin (**Figure 6A**, **Supplemental Figure 5A**). Strength of gene membership in the brown module negatively correlates with strength of correlation to serum insulin level, including *Fam151a*, which is highly connected in the brown network (**Figure 6B**). Further, expression of *Fam151a* negatively correlates with serum insulin levels across all cohorts, independent of network construction (**Figure 6C**). The brown network is visualized as the sum of gene-pair correlations (**Supplemental Figure 5C**). The network was subset into 20wk high-fat males (**Figure 6D**) and 20wk low-fat males (**Figure 5E**), revealing strong network structure in 20wk low-fat males. Differential connectivity was calculated between 20wk high and low-fat males to identify genetic network connectivity that was altered between cohorts (**Figure 6F, Table 14**). Differential connectivity was contrasted with differential expression between 20wk high and low-fat males, revealing *Fam151a* to be both differentially connected and differentially expressed (**Figure 6G**).

**Figure 6.**
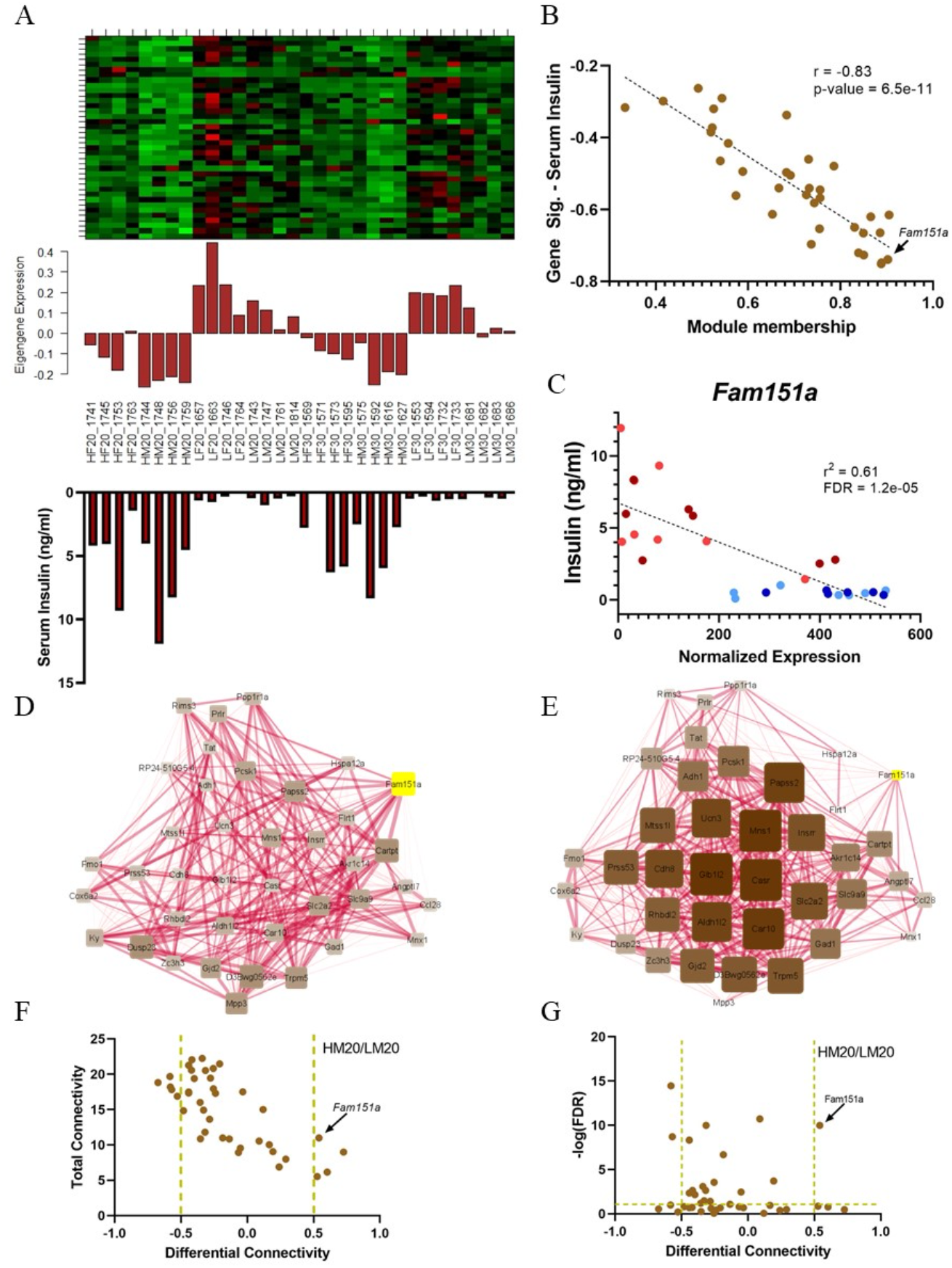
Brown module network structure is altered by diet in 20wk male mice. **(A)** Brown module heatmap, eigengene expression, and serum insulin levels across all individuals. **(B)** Correlation between strength of module membership and serum insulin levels for brown module. Gene of interest, *Fam151a*, highlighted as a highly connected member of the brown module and highly correlative with serum insulin levels. **(C)** Expression of *Fam151a* correlates with serum insulin levels across all individuals independent of module membership. Light red – 20wk high-fat mice, dark red – 30wk high-fat mice, light blue – 20wk low-fat mice, dark blue – 30wk low-fat fed mice. **(D)** Brown module network structure in 20wk high-fat males and **(E)** 20wk low-fat males. Gene of interest, *Fam151a*, is highlighted in yellow in each network. Size and color of node indicates overall connectivity within the network, thickness of edges indicates strength of correlation between gene pairs. **(F)** Total and differential connectivity between brown module genes in high and low-fat male SM/J mice. Vertical golden lines indicate threshold for significant differential connectivity. Gene of interest, *Fam151a*, is highlighted as a differentially connected gene. **(G)** Differential connectivity and differential expression of brown module genes between 20wk high and 20wk low-fat males. FDR – false discovery rate of p-values associated with differential expression. Vertical golden lines indicate threshold for differential connectivity, Horizontal golden line indicates threshold for differential expression. Gene of interest, *Fam151a*, is highlighted as differentially connected and differentially expressed.

### Single cell expression of Pdyn and Fam151a are influenced by age and diet

We sought to determine if the differential expression of *Pdyn* and *Fam151a* seen in the bulk analysis is recapitulated at the single cell level. We compared expression of *Pdyn* in all β-cells between 20wk high-fat males and 30wk high-fat males (**Figure 7A**, **Table 21**) and found significant differential expression, driven by differences in the proportion of β-cells expressing *Pdyn* (**Figure 7B**). Likewise, *Fam151a* is differentially expressed across all β-cells between 20wk high-fat males and 20wk low-fat males (**Figure 7C**, **Table 22**), driven by differences in the proportion of β-cells expressing *Fam151a* (**Figure 7D**). To determine if differential expression could be attributed to changes in β-cell subpopulation proportions, we calculated differential expression between the two most prominent subtypes, Beta 1 and Beta 2, revealing significant differential expression of *Pdyn* and *Fam151a* (**Figure 7E**, **Table 15**). While *Pdyn* is overexpressed in Beta 2 cells, which are prominent in 30wk high-fat males, *Fam151a* is overexpressed in Beta 1 cells, which are prominent in 20wk high-fat males. This suggests differential expression of *Pdyn* can be attributed to differences in subpopulation proportions, but not for differential expression of *Fam151a*. Assessing differential expression in Beta 1 cells between 20wk high-fat males and 20wk low-fat males revealed *Fam151a* was very significantly differently expressed (**Figure 7F**, **Table 23**), driven by an increase in the proportion of β-cells expressing *Fam151a* (**Figure 7G**), which could contribute to the differential expression of *Fam151a* between 20wk high-fat males and 20wk low-fat males seen in the bulk data.

**Figure 7.**
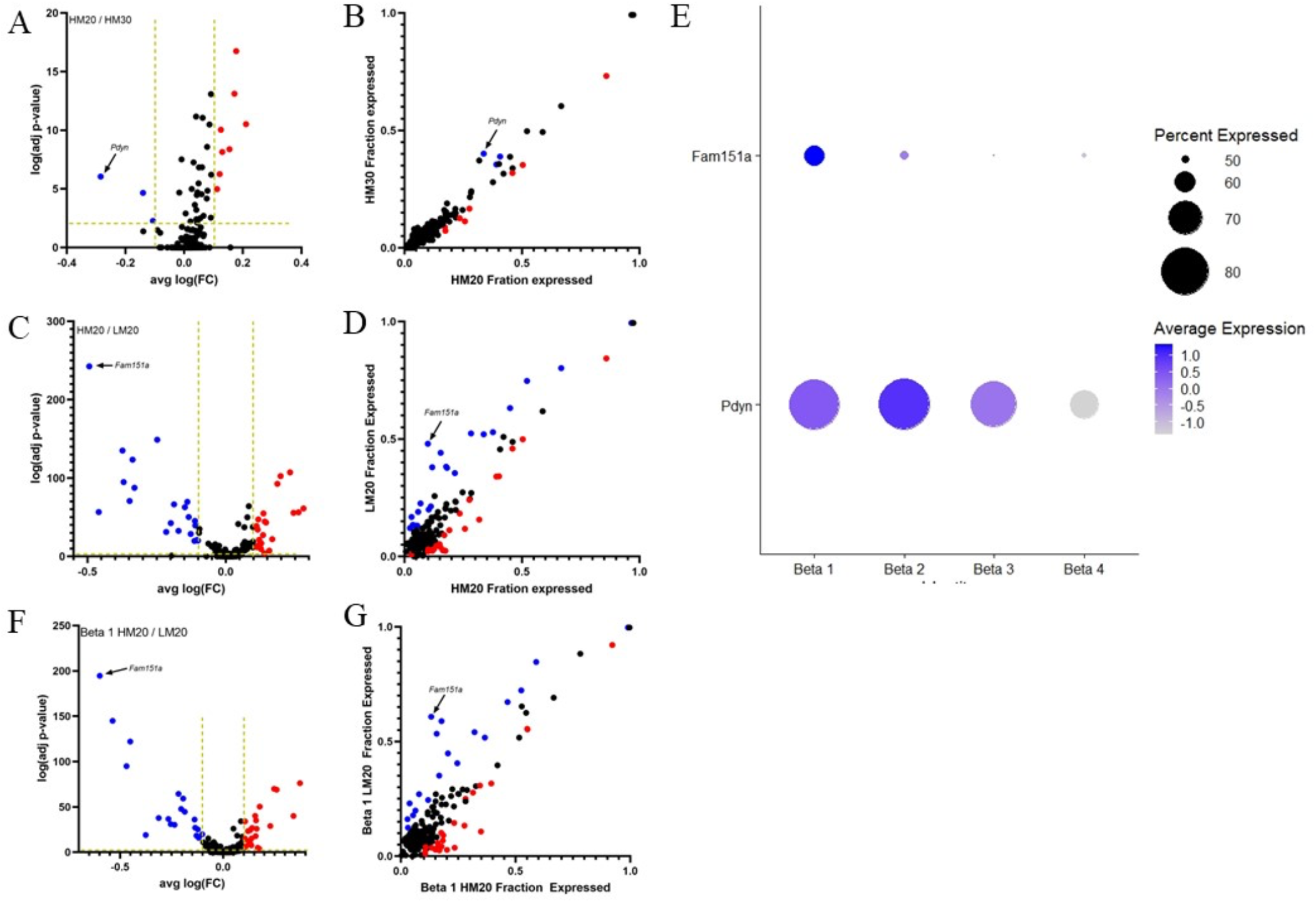
*Pdyn* and *Fam151a* are differentially expressed in β-cell subtypes. **(A)** Expression of β-cell-specific genes across all β-cells between 20wk and 30wk high-fat males. **(B)** Percent of β-cells in 20wk and 30wk high-fat males expressing β-cell-specific genes. **(C)** Expression of β-cell-specific genes across all β-cells between 20wk high and low-fat males. **(D)** Percent of β-cells in 20wk high and low-fat males expressing β-cell-specific genes. **(E)** Comparison of *Pdyn* and *Fam151a* expression across β-cell subtypes. **(F)** Expression of β-cell-specific genes in beta 1 cells between 20wk high-fat and 20wk low-fat fed mice. **(G)** Precent of beta 1 cells expressing beta specific genes between 20wk high-fat and 20wk low-fat fed mice. Blue genes are significantly under-expressed in comparison, red genes are significantly over-expressed. Vertical golden lines indicate threshold for significance based on average log fold change, horizontal line indicates threshold for significance based on Bonferroni corrected p-value.

## Discussion

Pancreatic β-cell heterogeneity has been studied extensively using single cell technology because of the cell type’s unique insulin-secreting capabilities and central role in diabetes etiology. Several groups have proposed the existence of functionally distinct β-cell subpopulations^35–38^, but their existence and relevance to insulin homeostasis remain controversial. We identified 4 β-cell populations (**Figure 1–2, Supplemental Figure 1**), Beta 1-4, across hyperglycemic-obese, normoglycemic-obese, and normoglycemic-lean mice. While the functional roles assigned to each population are based on gene enrichment analysis and open to interpretation (**Supplemental Table 2**), the relative proportion of each population changes in conjunction with islet function across cohorts, suggesting a functional relevance to overall β-cell population structure.

Beta 1 cells are most prevalent in hyperglycemic obese mice and are associated with elevated basal insulin secretion and low insulin content. Beta 1 cells overexpress mature markers including *Ucn3*^39^, *Pdx1*^40^, and *Acly*^41^, and negative regulators of glucose-stimulated insulin secretion including *Abcc8*^42^, *G6pc2*^43^, and *Ucp2*^44^. Terms associated with Beta 1 cells include signal release and hormone transport. Beta 1 cells appear to be primed for insulin release, but not to perform glucose-stimulated insulin secretion. Farack *et al*^36^ identified a subpopulation of “extreme” β-cells that specialize in basal insulin secretion, with high insulin expression and low mature insulin content. These cells have high expression of markers *Ucn3*, *Acly*, and *Pdx1*, and increase in proportion in obese mice. The Beta 1 cells we identified are similar to “extreme” β-cells based on their mature profile, potentially limited glucose response, and abundance in diabetic obese mice. Validation is required to determine if these β-cells function by over-expressing insulin secretion pathway components while suppressing glucose response mechanisms.

Beta 2 cells are prevalent in normoglycemic-obese and normoglycemic-lean mice, and are associated with high GSIS and high islet insulin content. Beta 2 cells are enriched for response to oxygen levels, pyruvate metabolism, and nucleotide phosphorylation, each associated with protection from hypoxia^45,46^. These cells are equally prevalent in obese 30wk high-fat males and lean 20wk low-fat males, which differ greatly in islet mass, suggesting Beta 2 cells are not a hypoxic population. Further, 20wk high-fat males have similar islet mass to 30wk high-fat males, but a much smaller Beta 2 population. Importantly, 30wk high-fat males have healthier glycemic parameters than 20wk high-fat males^20,21^. Several groups have identified a subset of heavily vascularized islets that have elevated oxygen consumption and superior GSIS at the cost of susceptibility to hypoxia. We suspect Beta 2 cells represent a highly functional β-cell population that confer protection against hypoxia^47,48^. Given their abundance in functional islets from both obese and lean normoglycemic mice, we hypothesize Beta 2 cells represent a mature, and possibly stress-resistant β-cell population.

Beta 3 cells are prevalent in normoglycemic obese mice which have elevated β-cell mass, high GSIS, and high insulin content. Beta 3 cells overexpress plasma membrane components, including several claudin family members, and ribosome components. Claudins provide structural integrity to tight junctions, which maintain cell polarity^49^. Polarity in β-cells serves as both a driver and a characteristic of mature, highly functional β-cells^35,50,51^. *Cldn4, Cldn3,* and *Cldn7* are all upregulated in Beta 3 cells and associated with mature β-cell function^52^. Beta 3 cells upregulate 7 ribosome genes and elevated ribosome biogenesis is associated with increased β-cell apoptosis^53^, β-cell replication^35^, and mature β-cell function^54^, making the functional relevance of overexpression unclear. One gene, *Rpl7*, is associated with mature β-cell function^55^, while *Rpl23* protects against apoptosis^56^. From this we conclude that Beta 3 cells represent a polarized and mature β-cell population.

Beta 4 cells are the lowest in abundance and elevated in lean males compared to obese males. Terms associated with Beta 4 cells include response to stress and response to topologically incorrect protein. In addition to overexpression of UPR components, Beta 4 cells have significant down regulation of *Ins1* and *Ucn3*. Several groups have identified β-cell subpopulations based on a stress response signature^13,57,58^, which have coalesced into a theory about stress response cycling, where β-cells go through periods of UPR activation and low insulin production to clear misfolded proteins^15,59^. Interestingly, 20wk low-fat mice have a similar proportion of stress response cycling cells compared to other groups (~7%), however this population is significantly smaller in high-fat males, suggesting that stress response cycling may be suppressed by obesity.

Efforts to describe β-cell transcriptional heterogeneity are marred by lack of consensus across scRNA-seq studies^60^. This is attributed to low read depth, which results in differences in gene capture driven by technical artifacts and random chance rather than biological variation^16^. Further, it is unclear what level of gene expression fold change is meaningful between individual cells, rendering single cell data poorly suited to assess differential expression across different cohorts. To address these problems, we developed a technique to integrate single cell data with bulk RNA-seq data (**Figure 3, Supplemental Figure 2**), which is not inhibited by these technical limitations, to assess β-cell-specific differential expression across cohorts of high- and low-fat fed male and female SM/J mice. While the list of genes identified as β-cell-specific is only 316 genes (**Supplemental Table 9**), we are highly confident of their specificity to β-cells, allowing for robust assessment of differential expression and network analysis. Sub-setting the bulk RNA-seq data by β-cell-specific genes provided a small but highly-focused search space to identify genes that influence β-cell function in hyperglycemic-obesity, normoglycemic-obesity, and healthy mice.

We identified *Pdyn* as a novel candidate gene associated with improved glycemic control in males. *Pdyn* encodes Proenkephalin-b, which is cleaved into Metenkephalin, an opioid with a short half-life and low bioavailability^61^. While *Pdyn*’s role in β-cell function is unknown, many opioids modulate insulin section^62^. Further, *Pdyn* expression is associated with regulation by *Pax6* and *Lkb1*, hallmarks of mature β-cells^63,64^. *Pdyn* is over expressed in Beta 2 cells, which are increased in normoglycemic-obese mice, and may link improved β-cell function with hypoxia response, as *Pdyn* provides protection from hypoxia in cardiomyocytes and neuronal tissue^65,66^.

We used weighted gene co-expression network analysis (WGCNA) to identify networks of co-expressed β-cell genes and to assess how networks correlate with metabolic and islet phenotypes (**Supplemental Figure 3**). Previous efforts to construct networks in islet RNA-seq data failed to account for cell-type specific gene expression, making it difficult to determine if these networks operate within an individual cell^67–69^. Our analysis revealed 6 β-cell-specific modules that correlate with phenotypes, two of which were enriched for intriguing ontology terms (**Supplemental Figure 4**). We chose to explore these networks in depth to assess the context in which they were relevant to β-cell function in obesity.

The blue module is highly expressed in 20wk high-fat males, strongly correlating with blood glucose concentration (**Figure 5, Supplemental Figure 5**). The blue module is enriched with fatty acid metabolism genes, which are heavily implicated in obesity and diabetes (**Supplemental Table 12**). In β-cells, fatty acid metabolism provides a glucose-independent stimulus for insulin release^70,71^. While short term fatty acid exposure enhances GSIS, prolonged exposure decreases GSIS and reduces insulin content^70,72^. Fatty acid metabolism protects against glucotoxicity-induced damage, allowing cells to convert excess glucose into glycerol, which bypasses ROS generation^73^. How this network connects to the β-cell dysfunction seen in obese, hyperglycemic 20wk high-fat mice is not immediately clear. Since fatty acid metabolism can stimulate insulin release, it is possible that prolonged activation of this network promotes elevated basal insulin secretion but inhibits GSIS. Expression of this network may be related to the abundance of Beta 1 cells in 20k high-fat males, which are geared toward basal insulin release. One gene in the blue module, *Entpd3*, confers susceptibility to diet-induced obesity and hyperglycemia^74^. This fatty acid metabolism gene network may provide a link between the increased proportion of Beta 1 cells and the β-cell dysfunction observed in 20wk high-fat males.

The brown module is highly expressed in lean mice and negatively correlates with serum insulin levels (**Figure 6, Supplemental Figure 5**). Unsurprisingly, this module is enriched for genes associated with β-cell physiology (**Supplemental Table 12**). We focused on how the network behaved in 20wk high and low-fat males, as they provided strong contrast for serum insulin levels and β-cell function. Core members of this network include known markers of β-cell maturation including *Ucn3*, *Slc2a2*, and *Pcsk1*. This network is suppressed in 20wk high-fat males, who have dysfunctional β-cells. We homed in on *Fam151a*, a gene highly connected within the network, that independently correlates with serum insulin levels. *Fam151a* is both differentially expressed and differentially connected within the brown module between 20wk high-fat and 20wk low-fat mice, suggesting an important role within the network and a potential link between serum insulin levels and β-cell function. *Fam151a* is a membrane embedded protein of unknown function, but it has been associated with dysregulated insulin secretion in *Isl1* knockout mice, and falls within a QTL associated with diabetes^75–77^. Further work will explore how *Fam151a* influences β-cell function in the context of obesity.

We explored if subpopulation composition contributes to differential expression of *Pdyn* and *Fam151a* (**Figure 7**). Total *Pdyn* expression increased across all β-cells between 20 and 30wk high-fat fed males, in agreement with the bulk RNA-seq data. *Pdyn* is overexpressed in Beta 2 cells, which increase in proportion between 20 and 30 weeks in high-fat mice, suggesting differential expression of *Pdyn* is driven by a change in subpopulation proportions. Investigation into how *Pdyn* influences in β-cell function must consider that knockout/overexpression may affect subpopulations differently, and that subpopulation structure changes based on metabolic context. Likewise, total *Fam151a* increased across all β-cells between 20wk high and low-fat males, in agreement with the bulk RNA-seq data. However, *Fam151a* is strongly overexpressed in Beta 1 cells, which are most abundant in 20wk high-fat males. To explore this discrepancy, we performed differential expression analysis in Beta 1 cells between 20wk high and 20wk low-fat males and found *Fam151a* expression is significantly higher in 20wk low-fat male Beta 1 cells. Thus, while the proportion of Beta 1 cells is lower in 20wk low-fat mice, the expression of *Fam151a* is significantly increased in their Beta 1 cells. These findings underscore the complexity of gene expression and suggest it is important to consider both subpopulation composition and expression within subpopulations when exploring β-cell heterogeneity in diabetes and obesity.

In summary, we identified 4 β-cell subpopulations whose relative proportions change depending on metabolic state. Beta 1 cells are primed for basal insulin secretion and proportionally high in hyperglycemic obese mice. Beta 2 cells are primed for protection from hypoxia associated with enhanced function and are abundant in normoglycemic obese and normoglycemic lean mice at the expense of Beta 1 cells. In conjunction, hyperglycemic obese mice express a highly connected genetic network associated with fatty acid metabolism, which is lost as glycemic control improves. The interplay between changing β-cell subpopulations and decreased fatty acid metabolism likely contributes to the improved β-cell function and subsequent restoration of glycemic control seen in obese SM/J mice^20,21^. This study provides a road map for exploring cellular heterogeneity by integrating sc- and bulk RNA-seq data, allowing for robust characterization of subpopulation structure, differential expression, and network analysis associated with obesity and glycemic stress.

## Author Contributions

HAL and MAM designed the experiments. MAM and HS performed mouse phenotyping and ELISAs. MAM and JFMV performed computational analysis and figure generation. MAM wrote the manuscript. All authors edited and approved the final draft.

## Funding

This work was supported by the Washington University Department of Genetics, the Diabetes Research Center at Washington University (grant P30DK020579), the NIH NIDDK (grant K01 DK095003 to HAL, grant T32 DK108742 to MAM, grant F31 DK125023-01 to MAM), GTAC@MGI Symposium Pilot Project Funding to HAL, and the NIH NHGRI (grant T32-GM007067 to JFMV).The authors declare no conflicts of interest.

## Table Legends

**Supplemental Table 1**. Differential gene expression between β-cell subtypes. Average log fold change (Avg_logFC) is calculated by dividing expression within subtype by expression in all other β-cells. Pct.1 is percent of cells in subtype expressing the gene, pct.2 is percent of all other β-cells expressing the gene. P_Val_adj is Bonferroni corrected p-value. Genes with Avg_logFC > 0.1 and p_val_adj <0.01 are considered significantly enriched in the subpopulation.

**Supplemental Table 2**. Type five results for over-representation analysis (ORA) significantly enriched genes within each β-cell subpopulation, including gene ontology terms.

**Supplemental Table 3**. Gene expression contribution for all cell types, in 20 week high-fat females.

**Supplemental Table 4**. Gene expression contribution for all cell types, in 30 week high-fat females.

**Supplemental Table 5**. Gene expression contribution for all cell types, in 20 week high-fat males.

**Supplemental Table 6**. Gene expression contribution for all cell types, in 30 week high-fat males.

**Supplemental Table 7**. Gene expression contribution for all cell types, in 20 week low-fat females.

**Supplemental Table 8**. Gene expression contribution for all cell types, in 20 week low-fat males.

**Supplemental Table 9.** β-cell-specific genes identified in single cell RNA sequencing analysis.

**Supplemental Table 10.** Top five results for over-representation analysis (ORA) for β-cell-specific genes, including gene ontology and mammalian phenotype ontology terms.

**Supplemental Table 11**. Bulk RNA sequencing differential expression analysis for β-cell-specific genes. For each analysis, log fold change, counts per million, p-value, and fdr corrected p-value are reported. hf = 20wk high-fat females/30wk high-fat females, hm = 20wk high-fat males/30wk high-fat males, lf = 20wk low-fat females/30wk low-fat females, lm = 20wk low-fat males/30wk low-fat males, f20 = 20wk high-fat females/20wk low-fat females, m20 = 20wk high-fat males/20wk low-fat males, f30 = 30wk high-fat females/30wk low-fat females, m30 = 30wk high-fat males/30wk low-fat males, age = all 20wk mice/all 30wk mice, diet = all high-fat mice/ all low-fat mice, sex = all females/all males.

**Supplemental Table 12.** Type five results for over-representation analysis (ORA) significantly enriched genes within brown and blue module genes, including gene ontology and KEGG pathway terms.

**Supplemental Table 13**. Gene significance and module membership for weighted gene co-expression analysis. For each gene, module membership is reported. Gene significance (GS) for each trait is reported as a Pearsons correlation, along with p-value (p.GS) associated with correlation. Module membership (MM) for each module reported as Pearsons coefficient along with p-value (p.MM) associated with correlation. BG = Blood glucose, GSIS= Glucose-stimulated insulin secretion, Basal.INS = Basal insulin secretion, INS.content = islet insulin content.

**Supplemental Table 14.** Network connectivity for all β-cell genes. Strength of connectivity for each gene within its assigned module reported as k-within value.

**Supplemental Table 15**. Differential gene expression between beta 1 and beta 2 subtypes. Average log fold change (Avg_logFC) is calculated by dividing expression within Beta 1 by expression in Beta 2. Pct.1 is percent of Beta 1 cells expressing the gene, pct.2 is percent of Beta 2 cells expressing the gene. P_Val_adj is Bonferroni corrected p-value. Genes with Avg_logFC > 0.1 and p_val_adj <0.01 are considered significantly differentially expressed.

**Supplemental Table 16**. Differential gene expression between beta 1 and beta 3 subtypes. Average log fold change (Avg_logFC) is calculated by dividing expression within Beta 1 by expression in Beta 3. Pct.1 is percent of Beta 1 cells expressing the gene, pct.2 is percent of Beta 3 cells expressing the gene. P_Val_adj is Bonferroni corrected p-value. Genes with Avg_logFC > 0.1 and p_val_adj <0.01 are considered significantly differentially expressed.

**Supplemental Table 17**. Differential gene expression between beta 1 and beta 4 subtypes. Average log fold change (Avg_logFC) is calculated by dividing expression within Beta 1 by expression in Beta 4. Pct.1 is percent of Beta 1 cells expressing the gene, pct.2 is percent of Beta 4 cells expressing the gene. P_Val_adj is Bonferroni corrected p-value. Genes with Avg_logFC > 0.1 and p_val_adj <0.01 are considered significantly differentially expressed.

**Supplemental Table 18**. Differential gene expression between beta 2 and beta 3 subtypes. Average log fold change (Avg_logFC) is calculated by dividing expression within Beta 2 by expression in Beta 3. Pct.1 is percent of Beta 2 cells expressing the gene, pct.2 is percent of Beta 3 cells expressing the gene. P_Val_adj is Bonferroni corrected p-value. Genes with Avg_logFC > 0.1 and p_val_adj <0.01 are considered significantly differentially expressed.

**Supplemental Table 19**. Differential gene expression between beta 2 and beta 4 subtypes. Average log fold change (Avg_logFC) is calculated by dividing expression within Beta 2 by expression in Beta 4. Pct.1 is percent of Beta 2 cells expressing the gene, pct.2 is percent of Beta 4 cells expressing the gene. P_Val_adj is Bonferroni corrected p-value. Genes with Avg_logFC > 0.1 and p_val_adj <0.01 are considered significantly differentially expressed.

**Supplemental Table 20**. Differential gene expression between beta 3 and beta 4 subtypes. Average log fold change (Avg_logFC) is calculated by dividing expression within Beta 3 by expression in Beta 4. Pct.1 is percent of Beta 3 cells expressing the gene, pct.2 is percent of Beta 4 cells expressing the gene. P_Val_adj is Bonferroni corrected p-value. Genes with Avg_logFC > 0.1 and p_val_adj <0.01 are considered significantly differentially expressed.

**Supplemental Table 21**. Differential gene expression between all β-cells in 20wk high-fat fed males and 30wk high-fat fed males. Average log fold change (Avg_logFC) is calculated by dividing expression within 20wk high-fat fed males by expression in 30wk high-fat fed males. Pct.1 is percent of 20wk high-fat fed male β-cells expressing the gene, pct.2 is percent of 30wk high-fat fed male β-cells expressing the gene. P_Val_adj is Bonferroni corrected p-value. Genes with Avg_logFC > 0.1 and p_val_adj <0.01 are considered significantly differentially expressed.

**Supplemental Table 22**. Differential gene expression between all β-cells in 20wk high-fat fed males and 20wk low-fat fed males. Average log fold change (Avg_logFC) is calculated by dividing expression within 20wk high-fat fed males by expression in 20wk low-fat fed males. Pct.1 is percent of 20wk high-fat fed male β-cells expressing the gene, pct.2 is percent of 20wk low-fat fed male β-cells expressing the gene. P_Val_adj is Bonferroni corrected p-value. Genes with Avg_logFC > 0.1 and p_val_adj <0.01 are considered significantly differentially expressed.

**Supplemental Table 23**. Differential gene expression between Beta 1 cells in 20wk high-fat fed males and 20wk low-fat fed males. Average log fold change (Avg_logFC) is calculated by dividing expression within beta 1 cells of 20wk high-fat fed males by expression in beta 1 cells 20wk low-fat fed males. Pct.1 is percent of Beta 1 cells in 20wk high-fat fed males expressing the gene, pct.2 is percent of Beta 1 cells in 20wk low-fat fed males expressing the gene. P_Val_adj is Bonferroni corrected p-value. Genes with Avg_logFC > 0.1 and p_val_adj <0.01 are considered significantly differentially expressed.

## Supplemental Figures

**Supplemental Figure 1.**
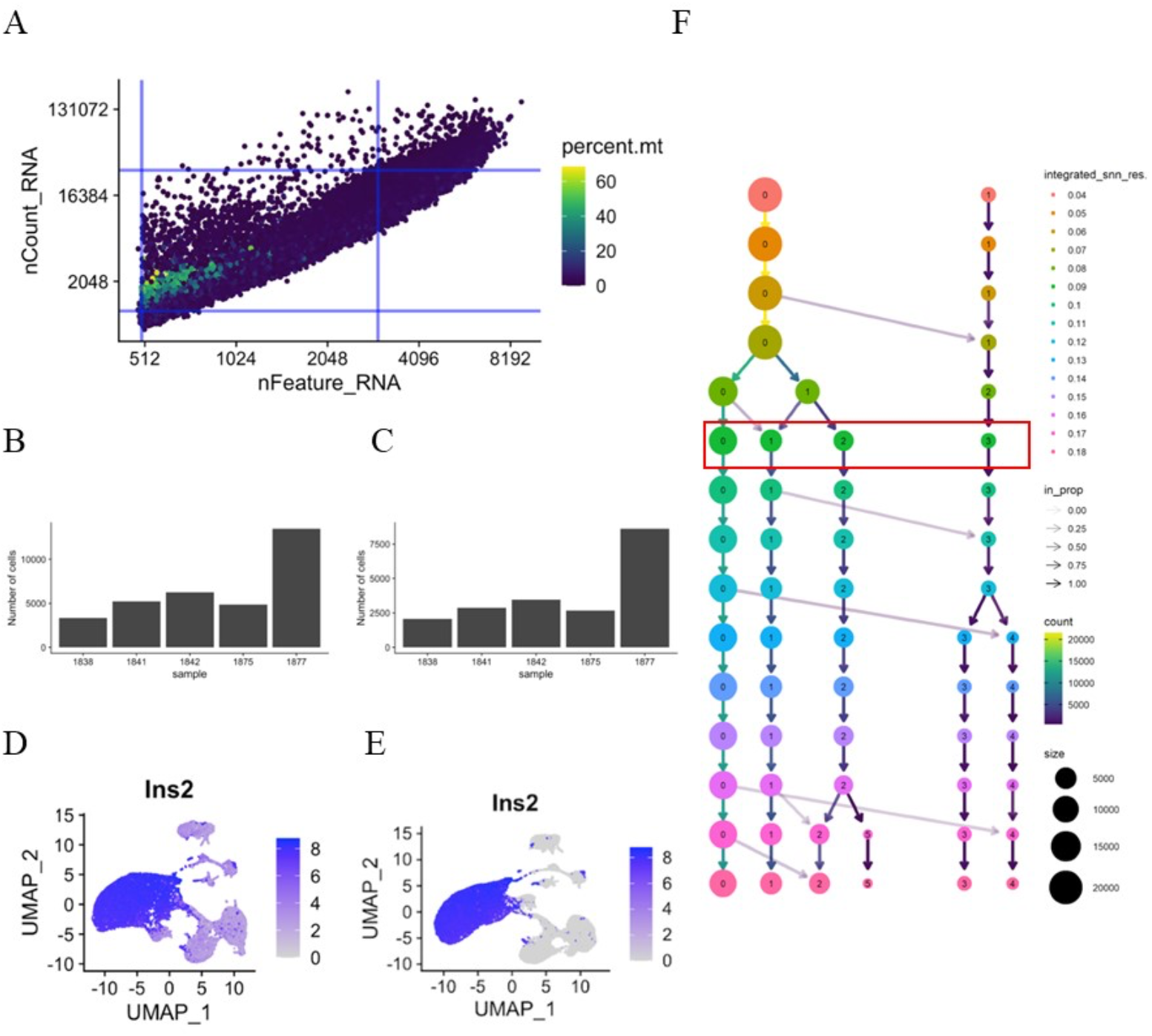
Single cell RNA sequencing quality control. **(A)** Total counts, features, and mitochondrial RNA expression plotted for each cell. Cells within blue box and without mitochondrial gene expression were included in analysis. **(B)** Number of cells identified in individual mice prior to quality control, **(C)** and after quality control. **(D)** Expression of *Ins2* across all cells prior to ambient RNA removing using SoupX, **(E)** and after ambient RNA removal. **(F)** Resolution analysis to determine the number of β-cell subpopulations. Selected resolution highlighted with red box.

**Supplemental Figure 2.**
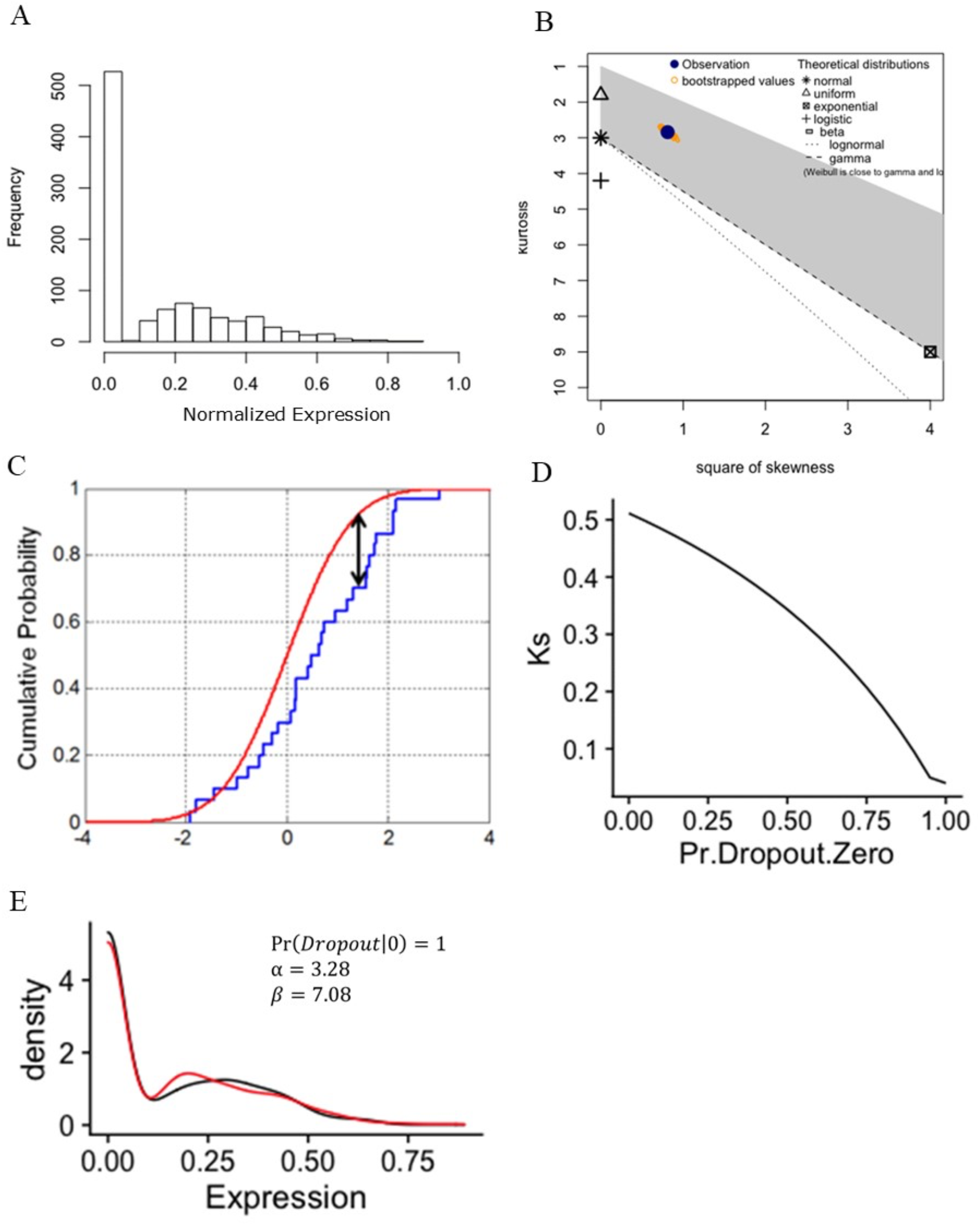
Quantifying expected expression for β-cell expression of *Dcd*. **(A)** Distribution of expression across all β-cells. **(B)** Cullen and Frey analysis identifies *Dcd* expression to be beta distributed. **(C)** Kolmogorov-Smirnov test identifies alpha and beta parameters that minimize Ks between real (blue line) and simulated (red line) distribution of cells expressing *Dcd*. **(D)** Estimating percent of cells not expressing *Dcd* due to gene drop out by iterating alpha and beta parameters that minimize Ks between real and simulated data. **(E)** Density plot visualizing estimated parameters for distribution of *Dcd* expression. Red line shows distribution of actual data, black line shows distribution of simulated data based on optimal alpha and beta parameters. From these parameters, expected expression is calculated.

**Supplemental Figure 3.**
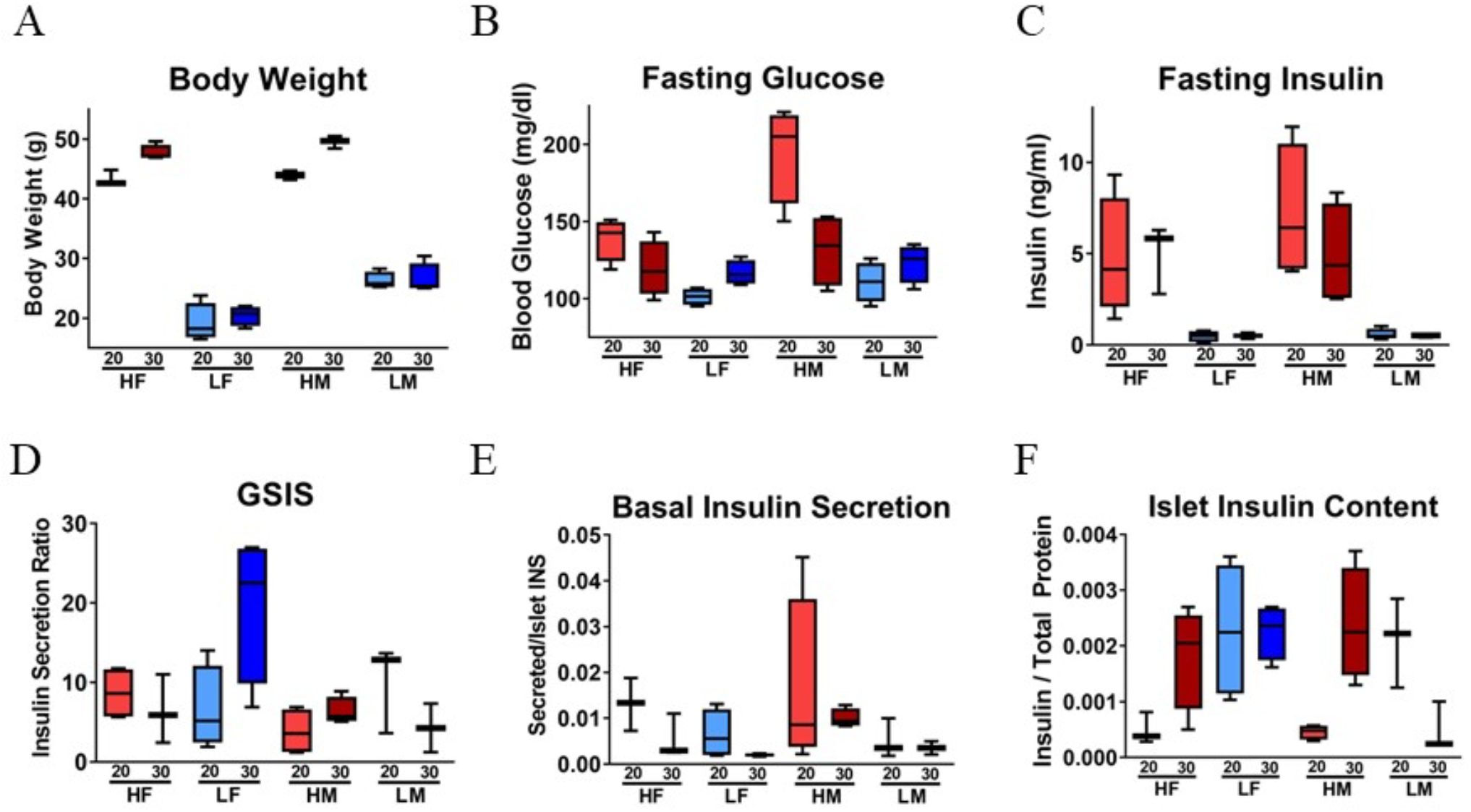
Metabolic and islet phenotypes from bulk RNA sequencing mice. **(A)** body weight, **(B)** blood glucose, **(C)** serum insulin levels collected after 4-hour fast. **(D)** Glucose-stimulated insulin secretion (GSIS), Basal insulin secretion, **(F)** Islet insulin content collected after isolated islets were rested overnight. N = 4 mice per age X sex X diet cohort. N= 10 islets per individual for islet phenotypes. Middle bar represents mean, box represents 25^th^ and 75^th^ quartile, whiskers represent minimum and maximum values.

**Supplemental Figure 4.**
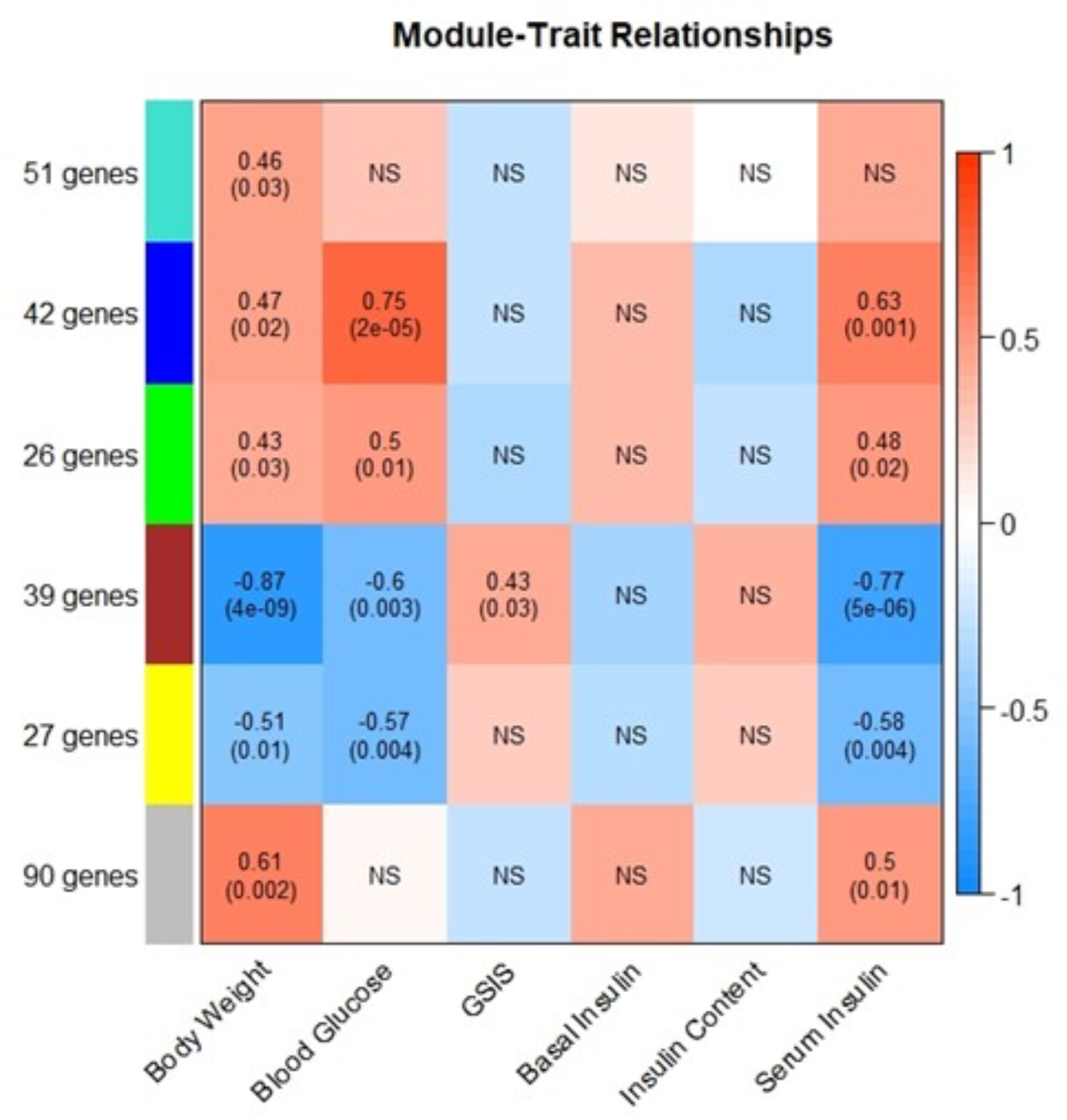
Correlation between phenotype and module eigengene expression. Number of genes within each module reported on y-axis, phenotypic trait reported on x-axis. For each module-trait relationship, the Pearson correlation between eigengene expression and phenotype value is reported (top) along with an FDR-corrected p-value for the correlation (bottom). Color of box indicates strength of correlation. NS = non-significant association based on FRD-corrected p-value.

**Supplemental Figure 5.**
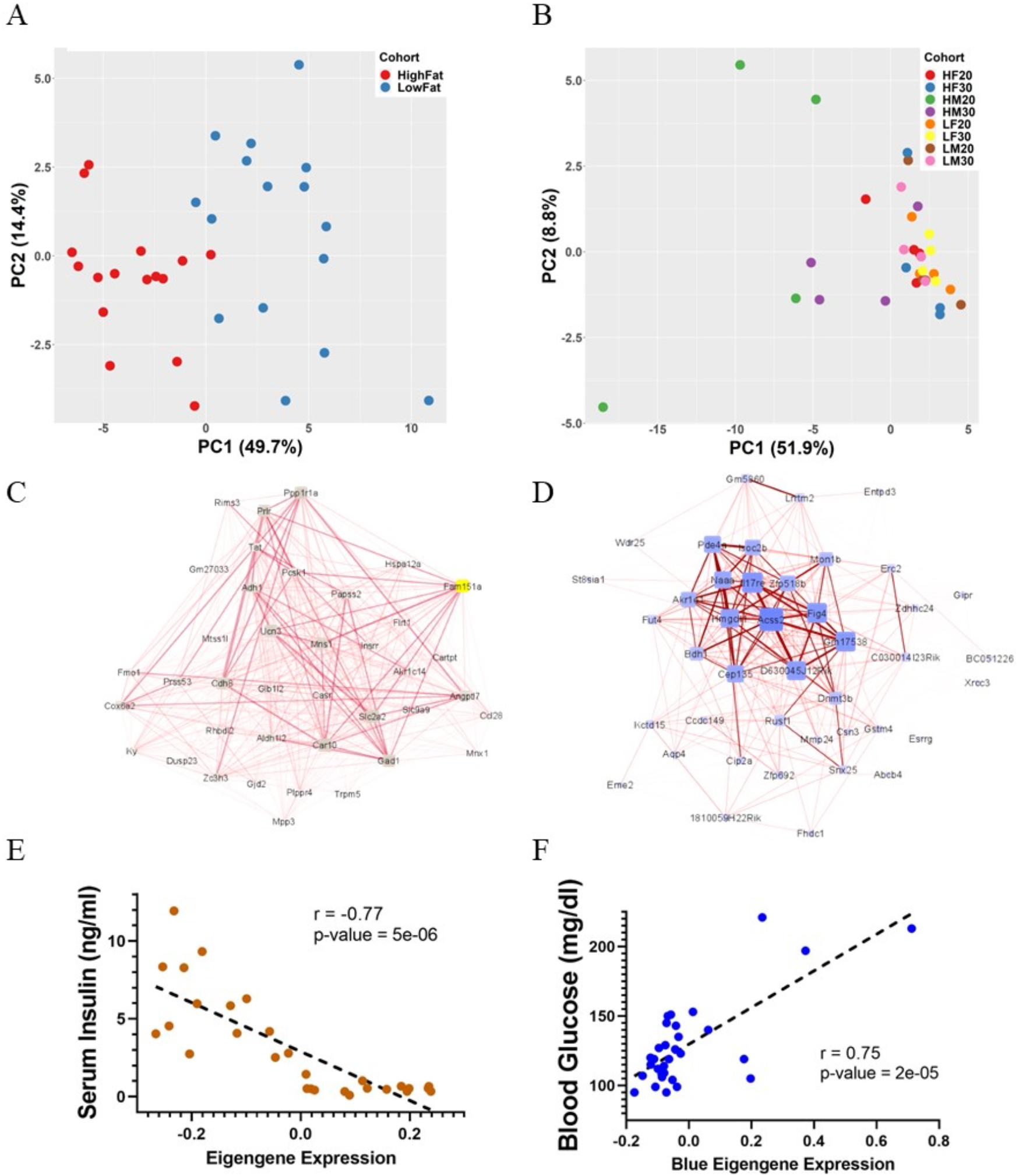
Overview of brown and blue modules. **(A)** Principal component analysis of gene expression within the brown module across all cohorts, segregated by diet. **(B)** Principal component analysis of gene expression within the blue module, segregated by cohort. **(C)** Brown module network structure in across all cohorts. Gene of interest, *Fam151a*, is highlighted in yellow. Size and color of node indicates overall connectivity within the network, thickness of edges indicates strength of correlation between gene pairs. **(D)** Blue module network structure in across all cohorts. Size and color of node indicates overall connectivity within the network, thickness of edges indicates strength of correlation between gene pairs. **(E)** Correlation between brown module eigengene expression and serum insulin levels across all cohorts. **(F)** Correlation between blue module eigengene expression and blood glucose levels across all cohorts.

## Notes

### Competing Interest Statement

The authors have declared no competing interest.

